# Severe Fe deficiency promotes hypoxia inducible responses in *Arabidopsis thaliana*

**DOI:** 10.1101/2025.01.07.631690

**Authors:** Yuri Telara, Moez Maghrebi, Mikel Lavilla, Noemi La Monaca, Giulia Ambrogini, Alessio Sbrana, Sara Delucchi, Pierdomenico Perata, Gianpiero Vigani, Beatrice Giuntoli

## Abstract

In plants, Fe homeostasis and O_2_ metabolism are strictly related, indeed several Fe-requiring enzymes catalyze reactions that also involve oxygen, as a reagent, product, entry or end point of the metabolic pathway in which the enzyme takes part. Oxygen sensing itself relies on Fe-dependent enzymes, the Plant cysteine oxidase (PCO) family of 2-OG independent thiol dioxygenases. PCOs are responsible for the degradation of ERFVII ethylene-responsive factors through a proteasomal N-degron pathway that connects hypoxia-inducible responses to the stabilization of the ERFVII transcription factors. Here, we investigated the interplay between low oxygen and Fe-deficiency stresses in *A. thaliana*. We used plants expressing a genetically encoded reporter of ERFVII protein stability and measured the expression of anaerobic genes to infer PCO activity *in vivo*. Our results highlight that Fe deprivation can elicit hypoxia-like responses depending on its severity. To test the involvement the ERFVII factors further, we examined the response of a pentuple *erfVII* mutant to Fe-deficiency stress, individually or combined with low oxygen. Our data indicate that the ERFVIIs might take part to the acclimation to chronic Fe deficiency by acting as positive regulators of starvation-responsive genes. Moreover, our results suggest that the ERFVIIs fine-tune nutrient mobilization to the shoots of submerged plants growing on moderately Fe-deficient substrates. This work expands the known functions of the ERFVII factors and provides new information to understand plant responses to combined environmental stresses.

## Introduction

Iron is an essential micronutrient for all living organisms, serving as a cofactor for numerous enzymes that regulate essential metabolic processes (Connorton et al., 2017; Vigani et al., 2013). Plants tightly regulate iron content to sustain growth and development, and the homeostasis of this micronutrient ultimately impacts on the quality and quantity of crop production (Kobayashi & Nishizawa, 2012). Iron supply limitations in agriculture are generally faced with the adoption of integrated soil and crop management practices, to prevent the occurrence in crops of nutritional disorders associated with Fe deficiency (Mahender et al., 2019.; Zuo & Zhang, 2011). Indeed, despite the abundance of iron in the Earth’s crust, its bioavailability is limited in nearly one-third of the world’s cultivated lands due to pH effects on soil redox state, which make iron less bioavailable to plants at neutral or alkaline pH (Manthey et al., 1994).

Fe uptake is enhanced through Strategy I, found in most flowering plants, or Strategy II mechanisms, restricted to grass species. Strategy I involves proteins like P-type H+-ATPase 2 (AHA2), FRO2, and Iron Regulated Transporter 1 (IRT1) genes, which work together to increase iron solubility and uptake (Morrissey & Guerinot, 2009). P-type ATPase release protons into the rhizosphere, reducing the pH and increasing Fe solubility. Ferrous oxidoreductases (FROs) exhibit Fe Chelate Reductase activity (FCR), converting Fe^3+^ into Fe^2+^. This FCR activity is enhanced at acidic pH (Tsai & Schmidt, 2017). Subsequently, ZIP transporters, such as IRT1 in *Arabidopsis thaliana*, facilitate the uptake of Fe^2+^ into plant roots. It has been demonstrated that AHA2, FRO2, and IRT1 form a protein complex at the root plasma membrane, which is suggested to create an optimal local environment with the right pH and Fe^2+^ concentration for efficient Fe uptake, preventing Fe^2+^ reoxidation in presence of oxygen in the soil (Martín-Barranco et al., 2020). Arabidopsis, a Strategy I plant, also shares characteristics with Strategy II by secreting Fe-mobilizing coumarins (FMC) from its roots into the rhizosphere. FMCs, derived from the phenylpropanoid metabolism, promote Fe uptake under conditions of low Fe availability (Robe et al., 2021).

The transcriptional regulation of Fe homeostasis in plants is a complex and critical process that governs proper acquisition, distribution, storage, and utilization of Fe while avoiding excessive accumulation that could lead to toxicity. In *Arabidopsis thaliana*, Fe homeostasis is regulated by two interconnected transcriptional modules. The first module involves the Fer-like Fe-deficiency Induced Transcription factor (FIT/bHLH29) from clade IIIe (Colangelo & Guerinot, 2004), which under Fe deficiency forms heterodimers with specific clade Ib TFs (bHLH38, bHLH39, bHLH100 and bHLH101) and activates the expression of *FRO2* and *IRT1* (Trofimov et al., 2019; Yuan et al., 2008; Wang et al., 2007) Conversely, clade IVa TFs (bHLH18, bHLH19, bHLH20, and bHLH25) repress FIT by promoting its degradation through the 26S proteasome pathway in a jasmonic acid-dependent manner (Cui et al., 2018). The E3 ligases Brutus Like 1 (BTSL1) and Brutus Like 2 (BTSL2) are additional negative regulators of FIT, again targeting it to the proteasome (Rodríguez-Celma et al., 2019). The second module involves the clade IVc bHLHs factors IAA-Leucine Resistant 3 (ILR3), Iron Deficiency Tolerant 1 (IDT1/bHLH34), bHLH104, and bHLH115 (Gao & Dubos, 2021), which play additive roles in the response to Fe deficiency. Their functions are partially explained by the ability to form homo- or heterodimers that induce *FIT* expression. Additionally, Iron Man/Fe-Uptake inducing peptides (IMA/FEP) participate to the Fe signaling cascade in Arabidopsis, with eight *IMA* genes regulating Fe homeostasis (Grillet et al., 2018). IMAs have been proposed to act as positive regulators of Fe uptake by regulating different components of Fe homeostatic control network. In particular, under Fe deficiency they can induce the expression of Ib clade bHLHs, *FRO2* and *IRT1* (Grillet et al., 2018; Hirayama et al., 2018). Also, IMA3 can inhibit BTS activity, buffering the degradation of ILR3 and bHLH115 proteins, and ultimately activating the Fe deficiency response(Li et al., 2021).

Such intricate regulatory mechanisms evolved to enable Fe homeostasis (preventing both Fe deficiency and toxicity) are linked to the central role that Fe chemistry plays in biological systems, established shortly after life appeared approximately 3.5 billion years ago when Fe(II) was abundant in the anoxic Earth environment (Ilbert & Bonnefoy, 2013). With the oxygenation of the Earth’s atmosphere, Fe chemistry and biochemistry have become heavily influenced by the presence of oxygen (Ilbert & Bonnefoy, 2013; Cammack et al., 1990). The functional relationship with oxygen can be incorporated in the classification of Fe-requiring enzymes (FeREs), along with their function (Vigani & Murgia, 2018a). With few exceptions, such as aconitase and purple acid phosphatase, FeREs carry out redox reactions that in many cases also involve oxygen (Cammack et al., 1990).

Dioxygenases are a wide class of enzymes that catalyze reactions where both atoms of molecular oxygen are incorporated into one or more substrates (Kawai et al., 2014; Kundu, 2012, 2015). Among the enzymes that use O_2_ as co-substrate, those belonging to this class are almost exclusively FeREs. Plant dioxygenases employ different types of Fe cofactors to activate oxygen in a variety of reactions with organic substrates, using different catalytic mechanisms (Iacopino & Licausi, 2020); (White & Flashman, 2016). The largest subfamily is formed by 2-OG/Fe-dependent dioxygenases (2-ODDs): their broad distribution in metabolism suggests that 2-ODD requirement for Fe may underlie the widespread effects that Fe deficiencies have in plants (Farrow & Facchini, 2014).

Among Fe-dependent dioxygenases, Plant cysteine oxidases (PCOs) are 2-OG independent enzymes that hold particular significance, as they function as genuine oxygen sensors (Gunawardana et al., 2022; Weits et al., 2014; White et al., 2018). PCOs affect the stability of group VII ethylene response factors (ERFVIIs), which are responsible for the fast activation of a core set of hypoxia-inducible genes conserved in plants (Abbas et al., 2015; Gasch et al., 2016). PCOs employ a mononuclear Fe(II) center to catalyze Cys-sulfinic acid formation at the N-terminal end of the ERFVII transcription factors, using O_2_ as co-substrate. Cysteine oxidation turns the ERFVIIs into targets for arginylation via ATE1/2, ubiquitination via the E3-ligase enzyme PRT6 and eventual degradation via the 26S proteasome (Gibbs et al., 2011; Licausi et al., 2011; White et al., 2017). Such proteolytic mechanism connecting protein half-life to O_2_ concentration is also known as cysteine or PCO branch of the N-degron pathway (hereafter, Cys-NDP). PCO inactivation under hypoxia blocks the Cys-NDP, leading to ERFVII accumulation and the induction of the anaerobic genes. Recently, a similar mechanism has been discovered in humans, where hypoxic responses are promoted by the homologous thiol dioxygenase ADO (Iacopino & Licausi, 2020; Masson et al., 2019).

Iron homeostasis and O_2_ metabolism are known to be linked in plants. Fe deficiency-induced responses are often associated with increased oxygen consumption rates in roots (Lopez-Millan et al., 2000; Vigani et al., 2009), whereas in leaves lower oxygen consumption and evolution have been described (Vigani et al., 2017). Despite these pieces of evidence, there is limited information regarding the specific impact of Fe deficiency on the dynamics of oxygen responses at the cellular level. The main goal of this study was to provide new insights in the interplay between Fe deficiency and hypoxic regulation in plants, stemming from the observation that O_2_ sensing relies on the Fe-requiring PCO enzymes.

## Materials and methods

### Plant material and growth conditions

The Columbia-0 ecotype (Col-0) of *Arabidopsis thaliana* was used as the wild-type background in all experiments. The pentuple mutant *erfVII* (Abbas et al., 2015) and the transgenic line *28RAPFluc (35S:RAP2.12_1-28_-FLUC)* (Weits et al., 2014) have been described previously. For axenic experiments, seeds of *A. thaliana* were surface sterilized with 70% (v/v) ethanol and 10% sodium hypochlorite and germinated on half-strength MS medium (Duchefa) supplemented with 10 g l^-1^ sucrose and 9 g l^-1^ agar. Plants were grown with 23°C day/19°C night temperature and a 12 h light period with a light intensity of 120 μmol photons m^-2^ s^-1^. Soil experiments were carried out according to (Murgia et al., 2015).

### Cloning of constructs

Plasmids pAG415-PCO4 and pAG415-GUS, used for yeast transformation, have been described before (Puerta et al., 2019).The chimeric DLOR-bHLH039 sequence, inspired to DLOR-RAP2.12 (Puerta et al., 2019) was designed by replacement of the original RAP2.12_2-28_ sequence with bHLH039_2-50_, and purchased as a synthetic Gateway^TM^-compatible (CACC-starting) DNA string from GeneArt (Thermo Fisher Scientific). The insert was cloned into pENTR-D/TOPO (Thermo Fisher Scientific) and subsequently recombined in the pAG413GPD-ccdB destination vector (Addgene plasmid #14142; Alberti et al., 2007) using the LR Clonase II Enzyme Mix (Thermo Fisher Scientific), according to the manufacturer’s recommendations.

The full length coding sequence of *bHLH039* was amplified from Arabidopsis cDNA, obtained as described in the paragraph “Gene expression analyses”. To amplify Gateway^TM^-compatible inserts with, respectively, Cys2 or Ala2 sequence, primers bHLH039(C)gw_FW (5’- CACCATGTGTGCATTAGTACCT-3’) or bHLH039(A)gw_FW (5’-CACCATGGCTGCATTAGTACCT-3’) were combined with primer bHLH039_RV (5’-TATATATGAGTTTCCACATTCC-3’). Amplifications were carried out with the high-fidelity Phusion DNA polymerase (Thermo Fisher Scientific), following the manufacturer’s recommendations. Fragments were cloned with the Gateway^TM^ system in the non-binary destination vector p2GWL7 (Weits, et al., 2014) obtained by ligation into the p2GW7 backbone of an ApaI/SpeI fragment excised from vector pBGWL7 both fromKarimi et al., (2002).

### Iron deficiency treatments

Nutrient composition of the modified MS media used for axenic experiments was described in detail by (Gruber et al., 2013). Fe-depletion treatments were performed by iron exclusion from the liquid media. Additionally, the agar was soaked in a solution containing 2 mM CaSO_4_ and 10 mM EDTA (pH 8) for 30 minutes and washed three times with pure H_2_O, each wash lasting 8 hours. Fe-chelation treatments were performed by supplementation of full nutrient media with 300 µM 2,2’-bipyridyl (Sigma-Aldrich) dissolved in DMSO (final DMSO concentration, 0.1% v/v); control samples here were treated with an equal dose of DMSO.

For short-term iron deficiency treatments, 7-day-old *28RAPFluc* seedlings were transferred from vertical plates to 6-well plates containing freshly prepared liquid media containing the specified iron concentration. Before the transfer, seedlings were kept in iron wash solution for 30 min, to remove iron from the apoplast, and rinsed with sterile distilled water.

In soil experiments, alkaline conditions were imposed according to Murgia et al. (2015). Control soil had pH 5.5, whereas alkaline soil (pH 7.8) was prepared by CaO supplementation (8 g CaO kg^-1^ substrate). The soil was moistened and thoroughly hand-mixed, its pH was measured after few hours and adjusted with further supplements of CaO, when needed. Soil was then mixed again and its pH measured several times in the next 2 days and each time adjusted to the target pH with CaO, before use. The pH was measured again at the end of the experiments, never exceeding 0.2 pH units drop.

### Low oxygen treatments

Hypoxic treatments were carried out in a Gloveless Anaerobic chamber (COY). Plants were treated in the dark with 1% O_2_ (v/v) atmosphere, or normoxic atmosphere, for the specified duration, at 23°C constant temperature. Dark submergence on 25 day-old soil grown plants was protracted for 12 hours, covering plants with 10 cm tap water column, pre-equilibrated at room temperature. Yeast cultures were treated for 6 h with 1% O_2_ (v/v) or normoxic atmosphere.

### Analysis of main root length

Primary root length of individual Arabidopsis seedlings was measured with ImageJ (Schneider et al., 2012). Plate images were taken at the end of the treatment period, using an Epson Expression 10000XL scanner (Seiko Epson) in color at 600 dpi resolution. Tests of statistical significance were undertaken with GraphPad 8.0 (Prism) using a two-way ANOVA with pairwise comparisons through the Tukey post-hoc test.

### Yeast culture and treatment

The heterologous assays were carried out in the *S. cerevisiae* strain BY4742 (*Matα; his3-Δ1; leu2-Δ0; lys2-Δ0; ura3-Δ0*) (Scientific Research and Development GmbH). Untransformed cells were grown on YPDA (20 g L^−1^ peptone, 10 g L^−1^ yeast extract, 20 g L^−1^ of glucose, 20 g L^−1^ agar, Duchefa). To generate the reporter strains, the pAG413-DLOR-bHLH039 plasmid was transformed along with pAG415-PCO4 or pAG415-GUS according to the LiAc/SS carrier DNA/PEG method, as previously described (Lavilla-Puerta et al., 2023). Transformed cells were grown at 30 °C in SD medium (6.7 g L ^−1^ Yeast Nitrogen Base DIFCO, 1.37 g L^−1^ Yeast Dropout Medium, 20 g L^−1^ glucose), plus adequate supplements (0.16 M uracil, 0.8 M histidine–HCl, 0.8 M leucine and 0.32 M tryptophan, when complete) and 20 g L^−1^ agar.

For the assay, five independent colonies were inoculated in 200 μl liquid SD medium with selection, in flat bottom 96-well polystyrene plates, in static regime. Overnight cultures were diluted to the early exponential phase with fresh media, allowed to resume growth and then adjusted to OD_600_=0.02 prior to treatment. The optical density of the cultures was measured directly in the microplate with a Multiskan Go 1510 Sky plate reader (Thermo Fisher Scientific). The cultures were then grown for 6 h under normoxic or hypoxic atmosphere, as illustrated above, with manual pipette mixing once in 2 h. At the end of the treatment, cells were recovered by centrifugation and subjected to luciferase assay.

### Protoplast transformation

Arabidopsis protoplasts were isolated from mesophyll cells and transfected as described in(Weits et al., 2014). In every independent transformation, 100 μl protoplast suspension, containing approximately 2x10^5^ cells, were co-transformed with 5 μg reporter plasmid (p2GW7-bHLH039 or p2GW7-RAP2.12_1-28_Fluc) and 2 μg p2GW7-Rluc normalizer plasmid. After 16 h incubation in the dark, protoplasts were recovered by gentle centrifugation and flash-frozen for the subsequent luciferase assays.

### Luciferase assays

Luciferase activity in *28RAPFluc* seedlings and in yeast microcultures was quantified using the Dual-Luciferase Reporter (DLR) Assay System (Promega) and a Lumat LB 9507 Tube Luminometer (Berthold), as described in (Lavilla-Puerta et al., 2023). For plant samples, values were normalized to the total protein amount, as determined through the Bradford protein assay (Bio-Rad), while for yeast samples values were expressed as Fluc/Rluc ratio.

### Immunoblotting

Total proteins were extracted from seven day-old Arabidopsis seedlings with 400 μL extraction buffer (50 mM Tris-HCl pH 7, 1 mM EDTA pH 8, 100 mM NaCl, 2% SDS, and 0.05% Tween-20 with a Protease Inhibitor Cocktail from Sigma). Extracts were centrifuged at 8000 g for 10 minutes at 4°C, and the protein concentration in the supernatant was measured using the Pierce BCA Protein Assay Kit (Thermo Fisher Scientific). 50 μg protein extract were denatured at 95°C for 5 minutes in 0.8 M DTT and XT Sample Buffer (Bio-Rad) and separated by SDS-PAGE on 10% polyacrylamide gels (NuPage Bis-Tris Gels, Thermo Fisher Scientific). Proteins were blotted onto a PVDF membrane (Bio-Rad) using the TransBlot® Turbo transfer system (Bio-Rad). For PCO1-GFP immunodetection, a monoclonal anti-GFP antibody (11814460001, Roche, 1:3000 dilution) was used and revealed with an anti-mouse IgG HRP (Agrisera AS09 627; 1:20000 dilution). An anti-HA peroxidase conjugated antibody (3F10, Roche, 1:1000 dilution) was used for the immunodetection of RAP2.3^3xHA^. Actin-11 was detected with a polyclonal primary antibody (AS13 2640, Agrisera, 1:5000 dilution) coupled with a goat anti-rabbit IgG HRP secondary antibody (Agrisera AS09 602; 1:15000 dilution). All antibodies were diluted in 4% skim milk solution in PBST. Signal detection was performed with Clarity Max Western ECL Substrate (Bio-Rad), using a ChemiDocTM MP Imaging System (Bio-Rad).

### Gene expression analyses

Total RNA was extracted as previously described (Kosmacz et al., 2015). RNA integrity was checked by gel electrophoresis on 1% (w/v) agarose, followed by spectrophotometric quantification. Reverse-transcription was performed with the Maxima First Strand complementary DNA (cDNA) Synthesis Kit (Thermo Fisher Scientific), following the manufacturer’s recommendations. RT-qPCR was performed with an ABI Prism 7300 sequence detection system (Applied Biosystems), using 12.5 ng cDNA template PowerUp SYBR® Green Master Mix (Thermo Fisher Scientific). *UBQ10* (*AT4G05320*) housekeeping gene expression was deployed for the calculation of relative gene expression following the ΔΔCt method (Livak & Schmittgen, 2001). Primer sequences are specified in **Supplemental Table 1**.

### ICP-MS measurements

Tissues (rosette leaves or entire seedlings) of plants were washed with Milli Q water and dried in a ventilated oven at 70 °C for 4 days. The dry weights were measured, and tissues were then digested with 500 µL 65% HNO_3_ for 4 h at 120 °C. Once mineralized, 250 µL of 65% HNO_3_ was added, and vortexed samples were transferred into polypropylene test tubes with a 1:40 dilution with Milli-Q water. Finally, the mineral contents of the samples were measured by inductively coupled plasma-mass spectrometry ICP-MS (BRUKER Aurora- M90 ICP-MS), as previously described (Vigani et al., 2021). The analyses were carried out from four independent biological replicates.

## Results

### Plant cysteine oxidase activity changes in iron-starved plants

Since PCOs are Fe-dependent enzymes, we speculated that they may function as convergence point between hypoxic signaling and Fe deficiency signaling. To investigate the effect of different Fe provisions on PCO activity, we deployed the *A. thaliana* reporter line *35S:RAP2.12_1-28_-Fluc*, expressing an ERFVII-based translational fusion (herafter indicated as 28RAPFluc) that informs about the activity of the Cys N-degron pathway (Lavilla-Puerta et al., 2023; Weits et al., 2014). In this line, firefly luciferase abundance is subjected to Cys2 regulation on the construct, but unaffected by mechanisms intervening on different domains of RAP2.12 beyond its N-terminus, such as signaling of the energy status (Weits et al., 2019) or proteolysis through SINAT1/2 E3 ligases (Papdi et al., 2015).

Preliminarily, we tested the responsivity of the reporter line to PCO inactivation by imposing hypoxia to 7 day-old seedlings, under Fe-replete conditions. In aerated samples, the signal remained unaltered over 18 h darkness, while it increased gradually in response to 3 and 6 h oxygen deprivation (dark hypoxia), reaching a maximum of 2.8-fold induction, with no further increase at 18 h **(Fig. 1a)**. We compared the output range under hypoxia, assumed as a strong inhibitory treatment of PCO activity, with 28RAPFluc response when Fe levels were manipulated under aerobic conditions. Complete removal of Fe from the apoplast and media, after supplementation of the Fe- chelating compound bipyridyl, caused up to 4.5-fold stabilization of the reporter **(Fig. 1b)**. The response had similar range to the one following acute hypoxia, but with faster and prolonged dynamics. ERFVII stabilization was accompanied by strong induction of downstream transcriptional responses, according to the anaerobic markers *Alcohol dehydrogenase 1* (*ADH1*)*, Lateral organ Binding Domain 41* (*LBD41*) and *PCO1* **(Fig. 1c)**. These observations suggest that plant PCOs can be readily and effectively inactivated by harsh Fe chelation treatments that make free Fe^2+^ unavailable. In parallel, we monitored the half-life of a GFP-tagged PCO1 version, stably expressed by *35S:PCO1:GFP* plants (Weits, et al., 2014) treated with the protein synthesis inhibitor cycloheximide (CHX). The immunoblotting showed that the fusion protein was characterized by long half-life **(Fig. 1d** and **Suppl. Fig. 1a)**. The slow turnover rate of PCO1 suggests that the iron cofactor was subtracted to previously formed PCOs, rather than made unavailable to newly synthesized enzymes.

**Figure 1.**
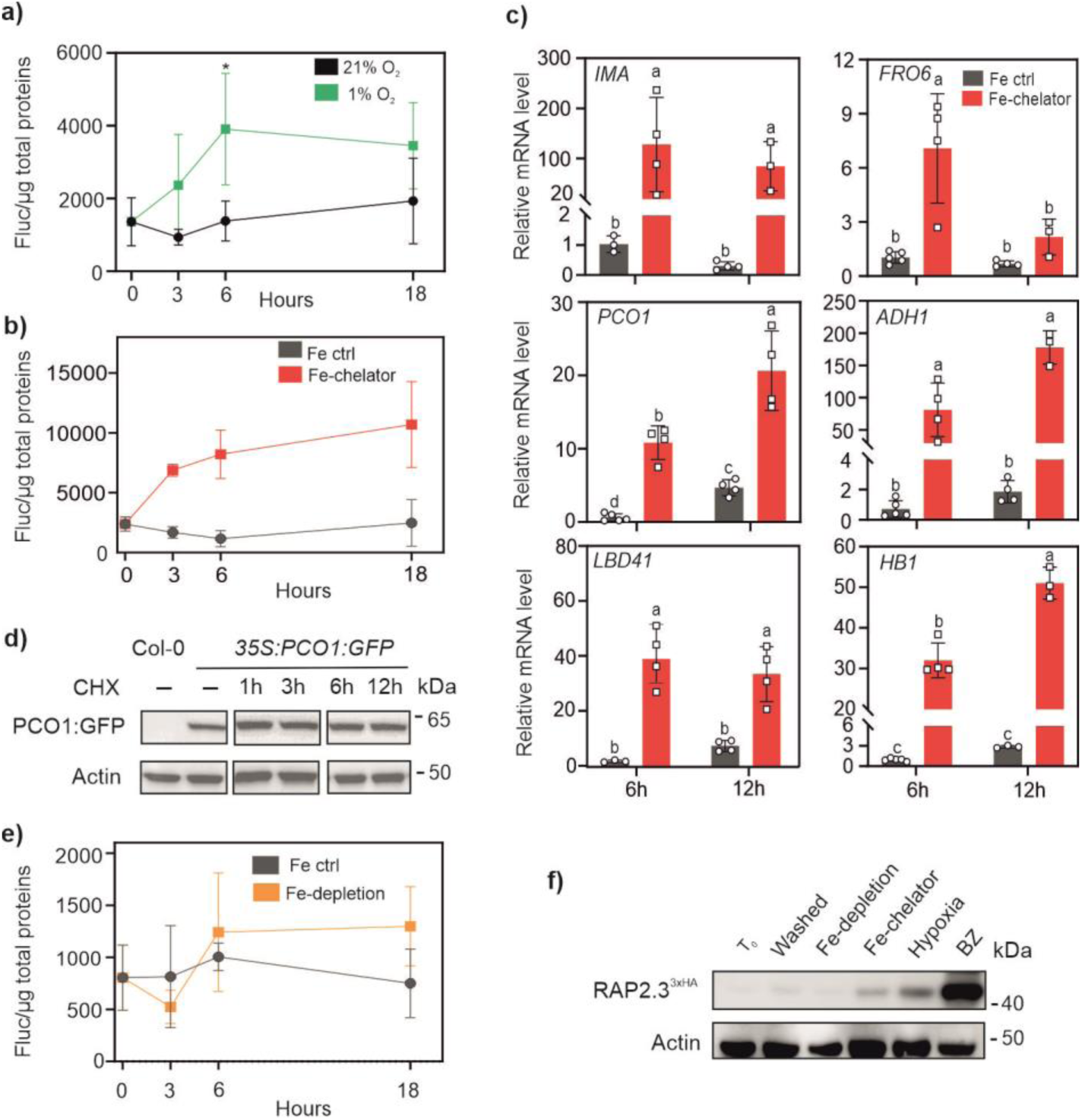
Modulation of anaerobic responses under different Fe-removal treatments. **(a)** Response of liquid grown five day-old *35S:RAP2.12_1-28_-Fluc* seedlings to hypoxia. Luciferase activity was measured at different time points of dark hypoxia (1% O_2_ v/v) or dark normoxia and normalized with soluble proteins (Fluc/µg total protein). Data in (a), (b) and (e) are mean ± SD (n=5). **(b)** Effect of Fe chelation on the 28RAPFluc reporter. Seedlings were transferred from vertical plates to 300 µM 2,2-dipyridyl (DIP), or to a control medium with 0.1% DMSO (v/v, “Fe-control”), and luciferase activity was measured after 3, 6 and 18 h treatment in the dark. **(c)** Anaerobic marker gene expression under Fe chelation conditions. Five day-old wild type seedlings, grown in liquid media, were treated with 300 µM DIP for 6 or 12 h in darkness, or with an equivalent amount of DMSO (Fe-control). The Fe deficiency-inducible gene *IRONMAN* was included to monitor the effects of iron chelation. Data are mean ± SD (n = 5). Letters indicate statistically significant difference (P< 0.05, two-way ANOVA, Tukey-Kramer post-hoc test). **(d)** Immunoblotting of *35S:GFP:PCO1* seedlings over a time course of cycloheximide treatment (CHX, 200 µM). **(e)** Effect of Fe deficiency on seven day-old *35S:RAP2.12_1-28_-Fluc* plants. Seedlings grown on full media in vertical plates were shifted to fresh Fe deficient (“Fe-depletion”) or full liquid medium (“Fe control”) after apoplast wash, and treated under neutral photoperiod. t_0_ corresponded to 4 p.m. Pairwise comparisons at every time point showed no significant differences (P>0.05). **(f)** Immunoblotting of RAP2.3 amount in *35S:RAP2.3^3xHA^*seedlings, seven day-old, grown in vertical plates and treated for 6 h in the dark. t_0_, before apoplast wash; washed, immediately after wash; Fe-depletion, Fe deficient medium; Fe-chelator, 300 µM 2,2-dipyridyl; hypoxia, 1% O_2_ (v/v) atmosphere; BZ, 300 µM bortezomib. ACT, actin-11 housekeeping protein. Supporting blots are provided in **Supplemental Figure 1**.

To simulate closer conditions to physiological Fe deficiencies, we imposed a milder treatment consisting in Fe removal from the apoplast by washing, followed by the transfer of seedlings on Fe- free media. In this case, Fe-depletion under photoperiodic conditions did not promote any stabilization of the 28RAPFluc reporter over 18 h treatment **(Fig. 1e)**. No reporter induction was observed when Fe depletion was combined with continuous darkness **(Suppl. Fig. 2a)**, ruling out that the exposure to light might interfere with the stabilization of the reporter in the previous experiment. We used the hemagglutinin-tagged line *35S:RAP2.3^3xHA^* (Gibbs et al., 2014)) to visualize ERF-VII abundance under iron deficiency by immunoblot. We could confirm that short-term Fe- depletion had no consequences on protein degradation, whereas Fe-chelation was comparable with 1% O_2_ (v/v) treatment in restraining RAP2.3 turnover **(Fig. 1f** and **Suppl. Fig. 1b)**. Proteasome inhibition after bortezomib (BZ) supplementation predictably had the strongest effect, due to fast and complete arrest of RAP2.3 degradation.

Longer Fe-depletion treatment, spanning over 48 h, was not able to promote 28RAPFluc stabilization either **(Suppl. Fig. 2b)**. In the same conditions, the anaerobic marker genes tested *PCO1, ADH1, PDC1* (*Pyruvate decarboxylase 1*) and *SAD6* (*Stearoyl-acyl carrier protein Δ9-desaturase 6*) showed sporadic and minimal responses to Fe-depletion, which were mostly recorded after the last time point **(Suppl. Fig. 2c)**. Such late induction appears to be unrelated to the early signaling events expected to follow PCO inactivation. Altogether, the previous observations indicated that transient Fe deprivation conditions do not generally associate with RAP2.12 stabilization. In contrast, Fe chelation achieved by bipyridyl treatments caused fast and significant stabilization of the ERFVII factors.

### Cys-NDP impact on the proteostasis of MC-containing Ib bHLH transcription factors

A potential hub between O_2_ and iron signaling is represented by the Fe-inducible subgroup of clade Ib bHLH transcription factors, composed of bHLH038, 039, 100 and 101 in *A. thaliana* (Wang et al., 2007), due to their conserved Cys2 residue **(Fig. 2a)**. Such observations led us to speculate of a functional role for Cys2 in making Fe-inducible clade Ib bHLH transcription factors susceptible to the Cys N-degron pathway.

**Figure 2.**
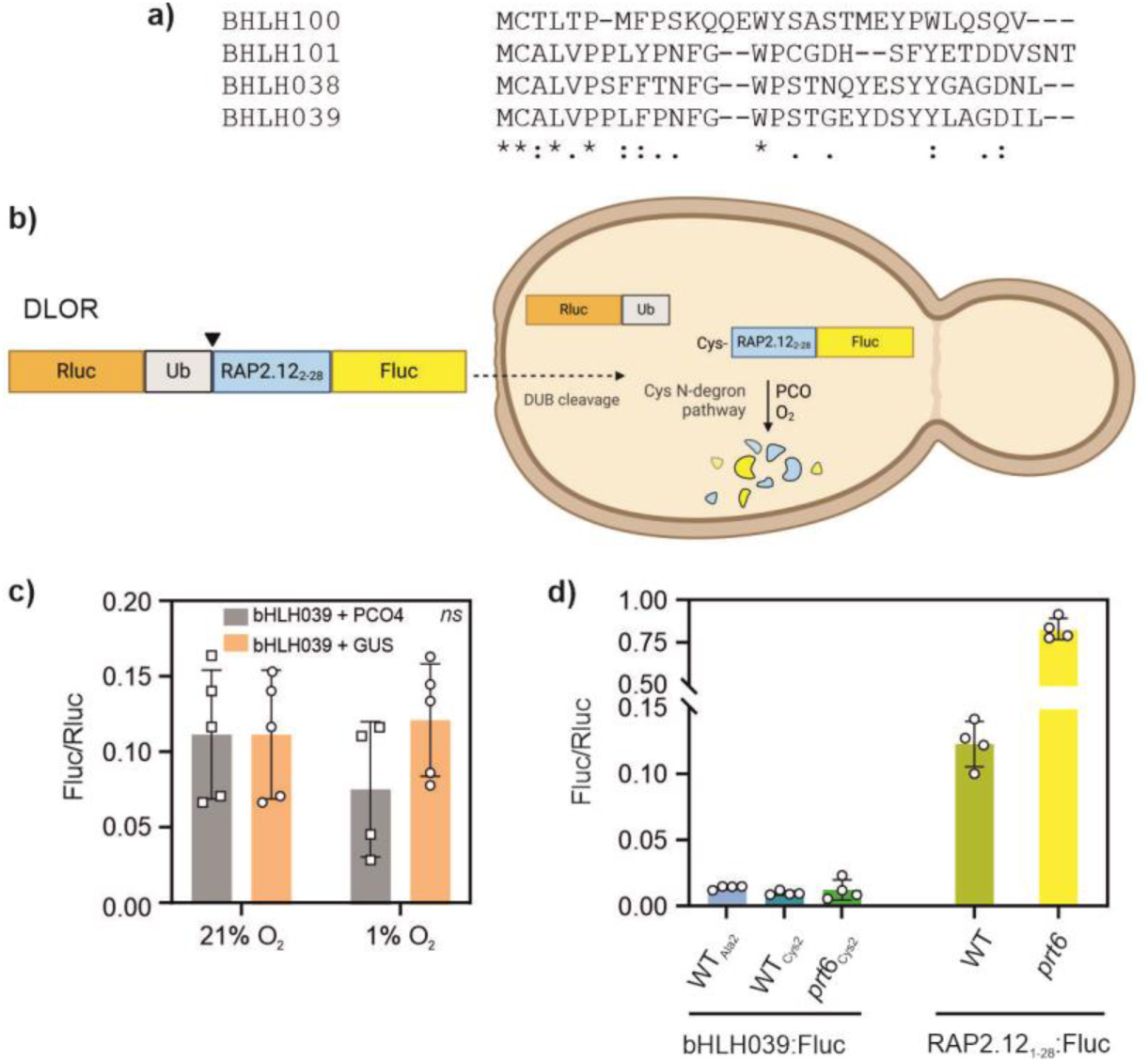
Subgroup Ib bHLHs turnover in relationship with the Cys-NDP. **(a)** N-terminal sequence alignment of the Cys2-containing subgroup Ib bHLH proteins of Arabidopsis. **(b)** Post-translational processing of the ratiometric reporter DLOR-bHLH039 in yeast. Once expressed in yeast cells, the chimeric protein is cleaved by native deubiquitinating enzymes (DUBs), which release a Cys-exposing fragment fused to firefly luciferase (Fluc) and the renilla luciferase protein (Rluc, used for normalization). Expression of heterologous PCO enzymes (here, PCO4 from *A. thaliana*) makes Cys- exposing substrates O_2_-labile through the native Arg N-degron pathway of yeast. Non-substrate proteins, instead, will remain stable. **(c)** Output of the DLOR-bHLH039 reporter, after 6 h treatment with hypoxia (1% O_2_ v/v) or normoxia (21% O_2_ v/v), in yeast cultures expressing AtPCO4 or a negative control construct (GUS, β-glucoronidase). Values are mean ± SD (n=4-5). Two-way ANOVA and Tukey-Kramer post-hoc test showed no significant differences (P>0.05). **(d)** Firefly luciferase activity of the bHLH039-Fluc reporter and the 28RAPFluc control construct (RAP2.12_1-28_-Fluc) expressed in Arabidopsis mesophyll protoplasts, normalized on total protein content in the extracts. Data are mean ± SD (n=4). WT, Col-0 wild-type. Cys2, wild-type *bHLH039* coding sequence; Ala2, Cys2Ala substituted version.

We chose bHLH039 as representative candidate from the set of four closely related proteins (Gao & Dubos, 2024). To evaluate its susceptibility to PCO-dependent degradation, we first resorted to an established reporter assay strategy deploying baker’s yeast (*Saccharomyces cerevisiae*) as heterologous testbed. We expressed a ratiometric luciferase-based reporter in yeast, DLOR-bHLH039, inspired to the one described in Puerta et al. (2019). DLOR-bHLH039 is a ubiquitin-fusion construct that, upon cleavage by endogenous deubiquitinating enzymes, releases a Cys-exposing fragment corresponding to the bHLH039_2-50_-Fluc sequence **(Fig. 2b)**. In this way, firefly luciferase (Fluc) activity provides an *in vivo* proxy of bHLH039 susceptibility to the N-degron pathway. In yeast cultures, DLOR-039 stability was unchanged regardless of AtPCO4 expression and atmospheric oxygen availability **(Fig. 2c)**. The substrate appeared to be very unstable in both tested conditions, hinting at PCO-independent degradation phenomena impinging on the protein fragment.

To exclude that the lack of regulation may be due to artefacts associated with yeast-specific regulation, we then evaluated bHLH039 stability in isolated mesophyll protoplasts of Arabidopsis. While the ERF-VII reporter construct 28RAPFluc was markedly stabilized in the *prt6* mutant (impaired in the last step of the Cys N-degron pathway), the full-length bHLH039-Fluc fusion protein was unaffected by either PRT6 inactivation or Cys2Ala substitution that would prevent PCO action **(Fig. 2d)**. Altogether, we could therefore rule out bHLH039 as a substrate of the Cys N-degron pathway. Moreover, the fusion protein emitted very low luminescence, suggesting that, in leaf cells, bHLH039 may be subjected to post-translational regulation to maintain protein levels low in Fe- replete conditions.

### Role of the ERFVII factors in the responses to prolonged Fe-deficiency

We showed before that the PCO branch of the N-degron pathway can be hampered in plants exposed to Fe-deficiency, leading to a reduction in ERFVII protein turnover **(Fig. 1)**. To delve more into the interplay between Cys-NDP and iron physiology, we examined the expression of ERFVII-regulated genes under chronic Fe starvation. Nutritional stresses were applied both to young Arabidopsis seedlings, germinating them on Fe-deficient axenic media, and rosette-stage plants, cultivated on alkaline soil (pH 7.8). Fe responses were monitored with the use of marker genes that are modulated by Fe deficiency regardless of plant tissues (*bHLH038, bHLH039, IMA*), along with *FRO* genes (Rodriguez Celma et al., 2019).

Wild type seedlings grown in plates displayed variable iron-deficiency symptoms across experiments. This outcome was not unexpected, since previous reports indicate that, for most elements, consistent levels of nutrient deficiency are difficult to reproduce when solid synthetic media are used (Gruber et al., 2013). The variability was likely due to persistence of traces of iron contaminant in the agar powder, after the agar washing procedure adopted. Although equal iron depletion over replicate experiments could not be achieved, this situation nonetheless gave us the chance to monitor plant responses under slightly different starvation conditions, at near-zero Fe levels.

In seedlings, permissive Fe deficiency was indicated by limited chlorosis, mild growth inhibition **(Suppl. Fig. 3a-b)** and moderate upregulation of Fe-deficiency markers in the wild type **(Suppl. Fig. 3c)**. Interestingly, no anaerobic gene induction took place in these conditions **(Suppl. Fig. 3c)**. The anaerobic response was also absent in rosette tissues of wild type plants cultivated on moderate alkaline soil for three weeks **(Fig. 3a)**. Supplementation of calcium oxide to the substrate increased soil pH from 5.5 to 7.8, causing a decrease of plant growth **(Fig. 3b)**. The small induction of the Fe- deficiency markers, along with weak chlorosis symptoms, suggested that these plants experienced moderate nutritional stress, similar to axenic seedlings above. This combined evidence suggests that plants affected by moderate Fe-deficiency for prolonged time did not generally invoke any long-lasting hypoxic signaling. We cannot rule out that transient hypoxic responses may have occurred at earlier growth stages, in starved seedlings or rosettes. However, our previous experiments with short-term treatments, failing to reveal any activation of low oxygen markers **(Fig. 1e and Suppl. Fig. 2)**, rather support the conclusion that hypoxic signaling is not elicited by mild Fe-starvation.

**Figure 3.**
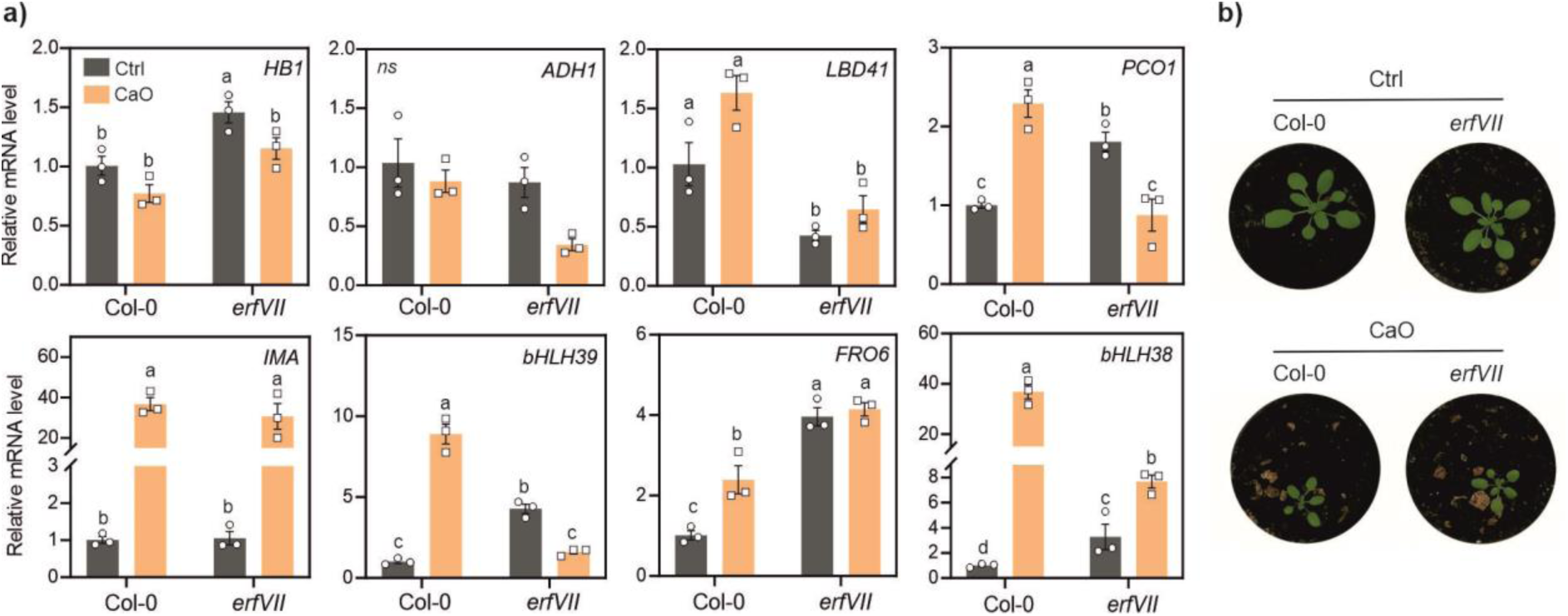
Response to chronic Fe deficiency in rosette stage plants. **(a)** Expression of anaerobic and Fe-starvation markers in rosette tissues of Col-0 wild type or *erfVII* mutant plants cultivated for three weeks on regular (Ctrl, pH=5.5) or alkaline substrate (+CaO, pH=7.8). **(b)** Representative pictures of plants used for the analysis.

Seedlings experiencing severe Fe deficiency displayed, instead, extremely stunted growth **(Fig. 4a, b)**, accompanied by strong upregulation of Fe-starvation markers still observable 10 days after germination **(Fig. 4c)**. Different from before, this harsh conditions strongly stimulated the induction of hypoxic markers **(Fig. 4c)**. The observed hypoxia-like response indicates that extreme chronic deprivation of Fe can lead to a prolonged impairment of PCO activity in plants.

**Figure 4.**
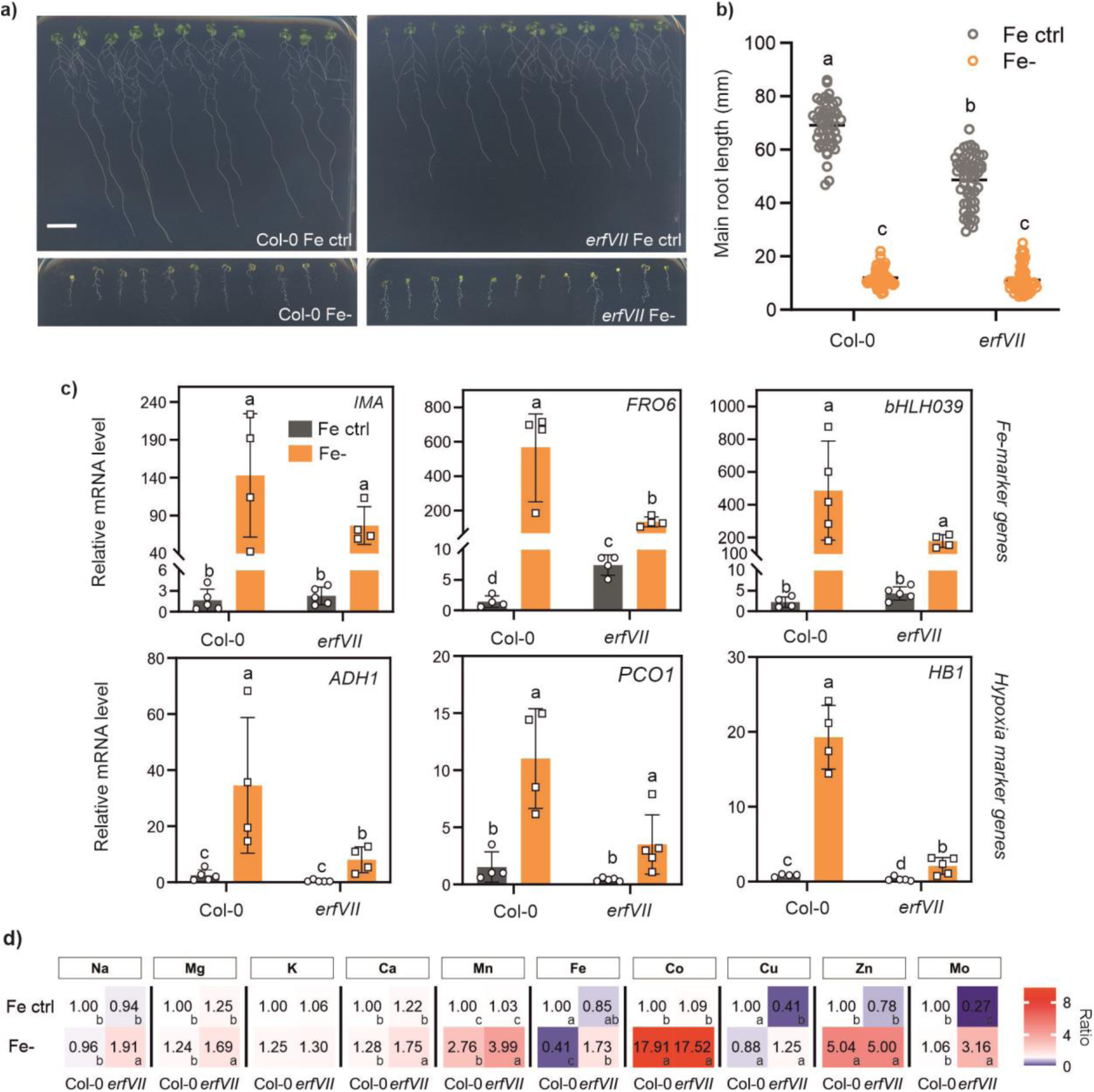
Response to chronic Fe deficiency in Arabidopsis seedlings. **(a)** Representative pictures of wild-type and *erf-VII* seedlings grown for 10 days on control or Fe- plates. Five replicate plates were observed for each experimental thesis. **(b)** Primary root length of plants in (a). Data are mean ± SD (n=57). **(c)** Expression of Fe deficiency- and hypoxia-markers in full seedlings from the same experiment. Expression values (mean ± SD, n=5) are presented as normalized to a wild type control sample. Distinct letters indicate statistically significant differences after two-way ANOVA and Tukey-Kramer post-hoc test (P< 0.05). **(d)** Heatmap of relative ion concentrations in seedlings, normalized on the average ion concentration in control Col-0 plants. Different letters indicate significant differences between conditions and lines (two-way ANOVA, Tukey’s test, P<0.05, n=3). Raw data are provided in **Supplemental Table 2**.

A pentuple *erfVII* mutant was included in the same experiments, to investigate how the modulation of the PCO pathway invoked by the stress may connect with physiological responses to Fe deficiency. In adult Fe-starved plants **(Fig. 3a)** and seedlings **(Fig. 4c)**, the upregulation of *bHLH039* was significantly lower in the *erfVII* mutant than the wild type. The consistent trend of transcriptional regulation of Fe- markers shown by plants at two distinct growth stages and from different substrates suggests that the ERFVIIs may take part to the acclimation to chronic Fe-deficiency overall plant life, by acting as positive regulators of starvation-responsive genes.

We evaluated whether the observed transcriptional regulation impacted on *erfVII* ability to cope with strong iron starvation at the seedling stage. After ten days on control media, the mutant produced significantly shorter primary roots than the wild type, in line with its previous phenotypic characterization (Shukla et al., 2019). When grown in strong Fe-deficiency conditions, instead, root elongation was not different between the mutant and the wild type **(Fig. 4a-b)**, ruling out a role for the ERFVII in primary root development under iron stress.

To gain insights in the physiological status of the treated seedlings, we determined the ionome profile of plants tissues by ICP-MS. We found no substantial differences in macronutrient content across genotypes and conditions, except for slightly higher levels of Ca and Mg in Fe- *erfVII* seedlings **(Fig. 4d** and **Supplemental Table 2**). On the contrary, Fe-deficiency markedly favoured micronutrient and metal ion (Co) intake, resulting in more elevated levels of three out of five micronutrients measured (Mn, Cu and Zn). No major differences between genotypes were again observed in Fe- starved plants, with the notable exception of Mo, whose homeostasis was completely altered in *erfVII*. The overall similarity in the nutrient profiles of the two genotypes was compatible with their comparable growth responses on Fe-. However, for some micronutrients (Mn, Cu, Mo) significant interaction between the variables was detected **(Fig. 4d** and **Supplemental Table 2)**. Therefore, while the ERF-VIIs did not regulate seedling growth in response to different Fe supplies, they are suggested to be involved in the fine-tuning of micronutrient content during chronic Fe deficiencies.

### Low-oxygen responses in control or Fe- conditions

Since the ERFVIIs were found to be connected with iron starvation responses, we decided to investigate the interplay between iron starvation and low oxygen conditions, where the ERFVII proteins are known to be stabilized and active. We asked ourselves whether iron availability may contribute to the full extent of anaerobic response in Arabidopsis plants and, on the other hand, whether the ERFVII factors may mediate Fe-starvation responses during hypoxia.

Wild type seedlings germinated as described before, on axenic Fe+ or Fe- media, were treated with short or prolonged hypoxia (1 or 6 hour treatment with 1% O_2_ atmosphere). Aerobic Fe- seedlings did not display any up-regulation of the anaerobic markers **(Fig. 5a)**, confirming that PCO catalysis was not affected by moderate Fe-depletion. Acute hypoxia, instead, triggered anaerobic gene expression, with up to 200-fold induction, which then progressed in a gene-specific fashion in the long term of stress. After 1 h, hypoxia marker transcripts were accumulated at the same levels in the two Fe supplementation regimes. This suggests that early marker gene induction, enabled by ERFVII stabilization, depended on quick PCO inactivation exclusively due to lack of sufficient oxygen for catalysis. After 6 h hypoxia, the effect of Fe-deficiency on anaerobic gene expression remained marginal: Fe-starved seedlings showed slightly less sustained expression of some markers (*HUP7, PCO1, LBD41*), while the other genes (*Hb1, PDC1, ADH*) were unaffected **(Fig. 5a)**. The data thus suggest that plant ability to respond to acute hypoxia stresses cannot be modulated by partial Fe removal from the environment.

**Figure 5.**
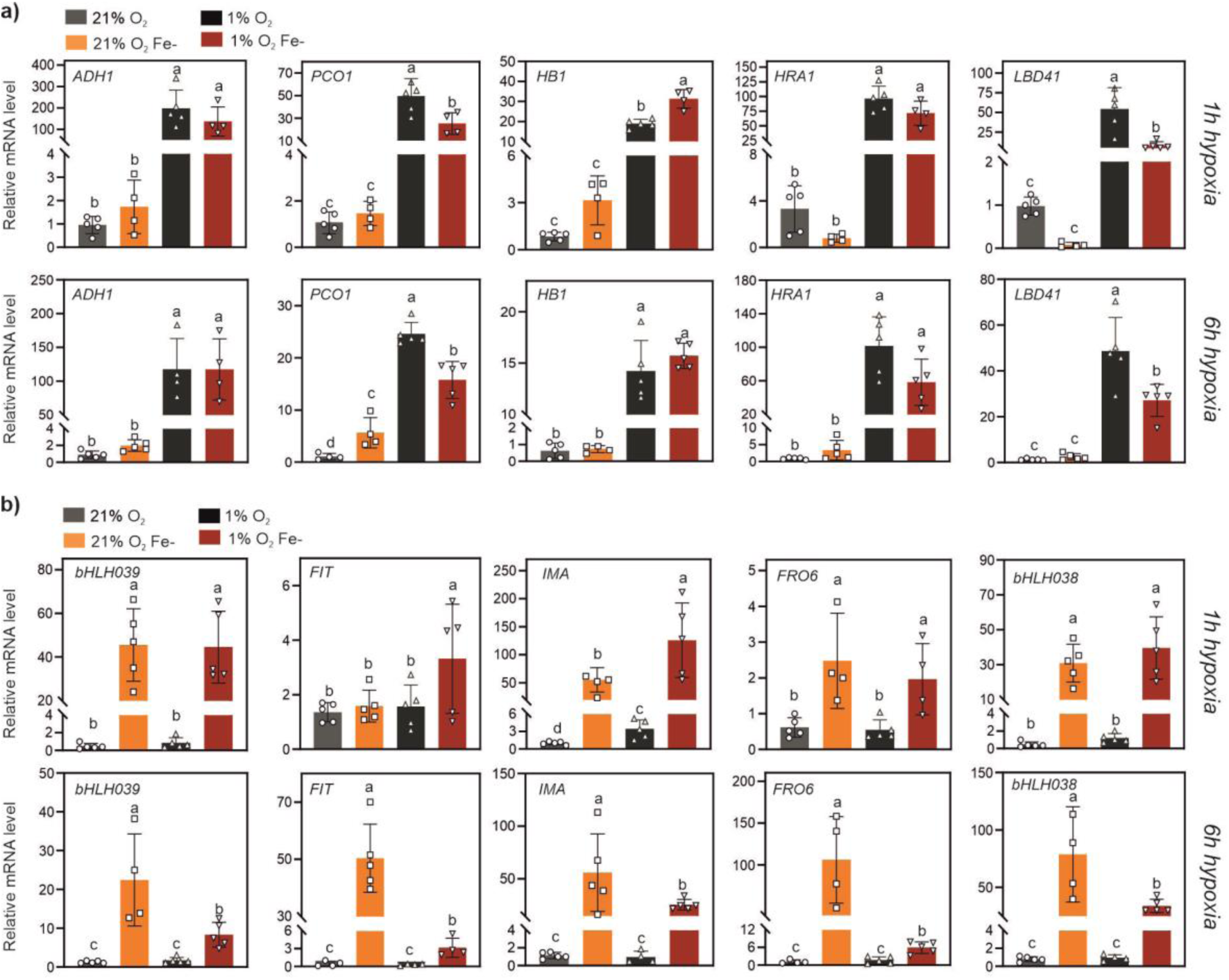
Effect of short and prolonged hypoxia on gene expression in seedlings grown under chronic iron deficiency. Expression of **(a)** anaerobic and **(b)** iron deficiency marker genes in Col-0 seedlings grown for 10 days under chronic iron deficiency and exposed to short hypoxia (1 h) or prolonged hypoxia (6 h) at 1% O_2_. Histograms show mean ± SD (n=5) of normalized expression values against one Fe-ctrl aerobic sample. Letters indicate statistically significant difference (P< 0.05, two-way ANOVA, Tukey-Kramer post hoc test).

We then looked into the regulation of five Fe-deficiency inducible genes during hypoxia **(Fig. 5b)**. In control plants, hypoxia did not promote the induction of any of the starvation marker tested, suggesting that ERFVII stabilization is not sufficient to promote Fe-deficiency responses. Furthermore, in Fe-starved seedlings (which as expected showed up-regulation of the markers, in normoxia), transcripts dropped when plants were shifted to hypoxia for 6 hours. This can be explained by selective transcription under hypoxia.

To reproduce a closer experimental setting to natural flooding combined with nutrient stress, we investigated the response of soil grown Arabidopsis plants to 12 hour submergence in presence of normal or alkaline substrate. We used ICP-MS to evaluate the impact of the interaction between iron starvation and hypoxia on nutrient acquisition. When wild type or *erfVII* plants were grown in normal soil, submergence had negligible effect on leaf ion content **(Fig. 6** and **Suppl. Table 3)**. Instead, a combination of submergence and alkaline conditions enhanced nutrient mobilization to the leaves (Ca, Mn, Mo and Fe itself) in the wild type, but not in the *erfVII*, as compared with control plants. We concluded that nutrient uptake during the first phases of submergence can be favoured by the ERFVII at high soil pH.

**Figure 6.**
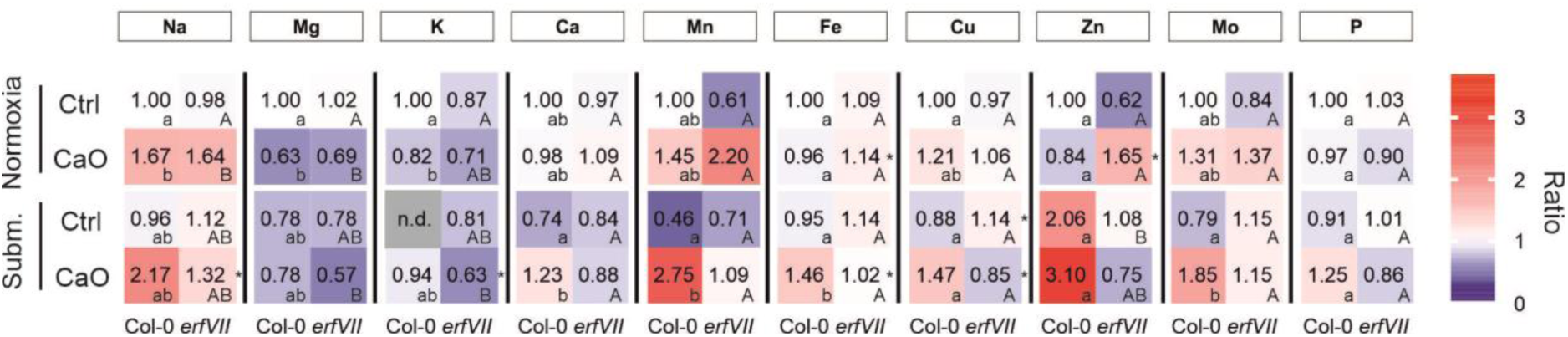
Heatmap of relative ion concentrations in Col-0 and *erfVII* plants treated or not with CaO and grown under normoxia and hypoxia conditions. Leaf content of inorganic elements from plants grown for three weeks in control (pH 5.5) or alkaline soil (pH7.8) (n=3). Ion abundance was expressed as the ratio to control Col-0 plants in normoxia and displayed as a heatmap. Ion concentration data are reported in **Supplemental Table 3.** Significant differences between the mean ion concentration of Col-0 and *erfVII* (within conditions) are indicated by an asterisk (P<0.05 after t-test). Different letters indicate significant differences among conditions within genotypes, after two-way ANOVA followed by Tukey’s test (P<0.05). Lower and upper case letters were used for Col-0 and *erfVII*, respectively.

CaO supplementation resulted in a moderate alkaline condition (pH 7.8), which, under normoxia, did not alter the total Fe content of leaves in 21-days old plants **(Fig. 6)**, which are prioritized tissues in terms of iron transport (Vigani et al., 2017). Nonetheless, a transcriptional analysis revealed that Fe- responsive markers except for *FIT* were induced in leaves from the same plants **(Fig. 7a)**, indicating the activation of Fe deficiency-induced responses. Soil alkalinization was not sufficient to elicit any hypoxia-like response in normoxic plants, as expected from the previous observations, nor it impacted on anaerobic marker gene upregulation by submergence **(Fig. 7b)**. This suggests that submerged plants growing on moderately alkaline substrates are able to mount normal anaerobic responses (as inferred from core hypoxia-inducible genes).

**Figure 7.**
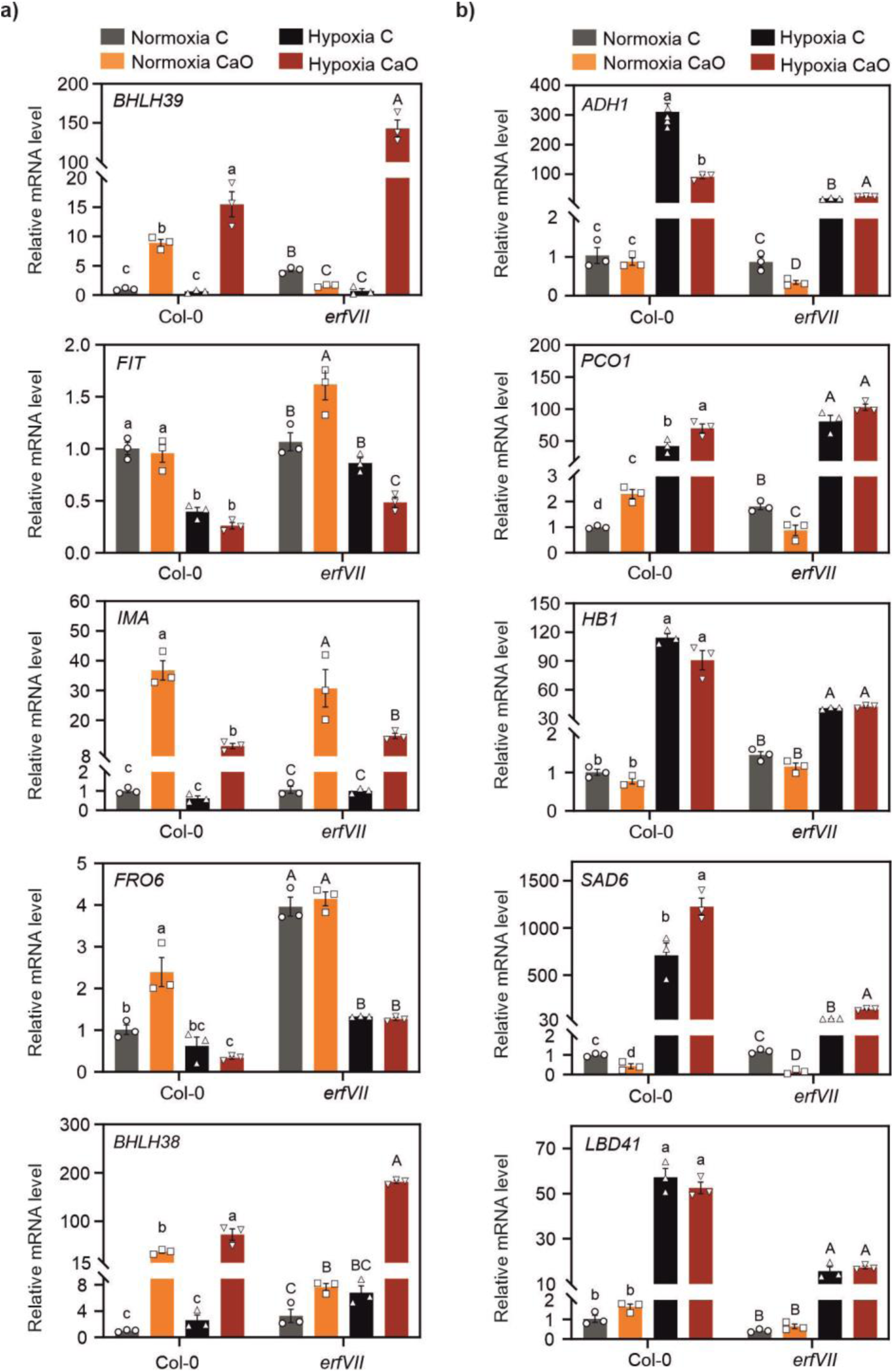
Molecular responses in *erfVII* mutants grown in alkaline soil. Expression of **(a)** Fe deficiency and **(b)** anaerobic genes in leaves of Col-0 and *erfVII* plants grown for three weeks in control (pH 5.5) or alkaline soil (pH 7.8) (n=5). Letters indicate statistically significant differences within the same genotype (P< 0.05, two-way ANOVA, Tukey-Kramer post hoc test).

Submergence *per se* did not affect the expression of Fe-deficiency markers **(Fig. 7a)**, similar to hypoxia in seedlings **(Fig. 5)**. However, submergence combined with alkaline soil conditions put *erfVII* mutant plants into a mild Fe starvation state, not experienced by the wild type. Among the Fe marker genes that displayed significant variations in leaves, *bHLH038* and *039* (which under normoxia had lower expression than in the wild type, as already shown in **Figure 3**) were indeed more expressed in *erfVII* than Col-0 plants grown on CaO during submergence **(Fig. 7a)**. *IMA* expression, instead, was comparable to the wild type. The stimulation of Fe-deficiency responses under stress was in line with lower micronutrient mobilization capacity in the mutant. Moreover, the favoured differential behaviour of the tested markers suggests that the ERFVII factors are involved in the regulation of a subset of Fe responsive genes, by restraining their induction in submerged leaves of Fe starved plants.

## Discussion

Plants have evolved sophisticated mechanisms to sense and respond to environmental fluctuations, leading to an increase in the frequency and distribution of different stress factors worldwide. The co-occurrence of multiple stresses may negatively impact plants on growth and health through synergistic or antagonistic interactions among different pathways, networks, and mechanisms that are promoted by each of the stress factors (Zandalinas & Mittler, 2022). Here, we examined the molecular cross-talks between Fe-deficiency and low oxygen response pathways in the model plant *A. thaliana*, which have received little attention so far.

Fe and hypoxia are expected to be interconnected at two major points in plant physiology: 1) the phytohormone ethylene is a mediator of hypoxia and Fe deficiency responses (Li & Lan, 2017; Lucena et al., 2015), and 2) non heme Fe-dependent dioxygenases are responsible for oxygen sensing (Perri & Licausi, 2024; White et al., 2017, 2018). In our work, we moved from the latter, asking ourselves whether PCOs might serve as multisensors of Fe^2+^ and O_2_ fluctuations. A main objective of this study was therefore to investigate the sensitivity of PCO enzymes to Fe deprivation, in physiological Fe-deficiency situations. Several studies indicate that Fe economy in plants involves the differential regulation of the various iron-requiring enzymes (FeREs) at different stages of severity of Fe deficiency stresses (Blaby-Haas & Merchant, 2013;Vigani & Murgia, 2018). The preferential use of Fe by specific metabolic pathways under Fe deficiency, known as “priority of Fe use”, is believed to serve as an acclimation mechanism to preserve Fe for the most important functions (Hantzis et al., 2018) and represents a hallmark of the phase of resistance to the stress (Vigani & Murgia, 2018). By evaluating ERFVII stability and the expression of anaerobic marker genes as proxies of PCO activity **(Fig. 1** and **4)**, we could conclude that PCO catalysis is only affected when Fe is left in trace amounts in tissues. This suggests that PCOs might be identified as prioritized components during Fe deficiencies, meaning that PCOs engagement might occur later in the progression of Fe deficiency induced responses.

The differential engagement of FeREs can be explained by their affinity for Fe as compared with the physiological concentration of free redox-active Fe ions, also known as Labile Iron Pool or LIP. FeREs characterized by K_mFe(II)_ values close to LIP concentration at a specific subcellular location will be strongly affected by Fe fluctuations, becoming partially or completely inhibited as Fe availability falls below the physiological LIP. In contrast, FeRE with lower K_mFe(II)_ than the physiological LIP may remain active even under Fe deficiency (Vigani et al., 2013). The former behaviour is expected to be associated with Fe sensing functions. Based on *in silico* predictions, it has been proposed that the plant 2-OG dependent dioxygenase (2-ODD) superfamily of O_2_-dependent FeREs might harbour enzymes with iron sensing potential (Kundu, 2015a; Vigani et al., 2013). However, such a role has been ruled out for the human prolyl-4-hydroxylases (PHDs), 2-ODD enzymes that serve as oxygen sensors in metazoans. In fact, it has been ascertained that PHD2 can form stable complexes with Fe(II) (Flashman et al., 2010; Hirsilä et al., 2003).

No experimental data of K_mFe(II)_ are available, instead, for thiol dioxygenases. It has been shown that excess of divalent Zn ions can displace Fe(II) from the catalytic site of PCOs and, thereby, lead to the activation of Fe deficiency-like responses in poplar and Arabidopsis (Carbonare et al., 2019a). Purified PCO and cysteine dioxygenase enzyme (CDO, a closely related dioxygenase present in metazoans) have been suggested to be unable to bind Fe(II) tightly, based on their binding of substechiometric amounts of the cofactor (Imsand et al., 2012; White et al., 2018). However, exogenous Fe(II) supplementation to the five different purified PCO isoforms of Arabidopsis enhanced PCO2 specific activity, but had no effect on the other isoforms. In our *in vivo* Fe-removal experiments, Fe withdrawal from the media had no effect on reporter stability over 18 hours, unless a chelator was added. In parallel, we showed that PCO1 had an observed half-life above 12 hours in Arabidopsis seedling tissues **(Fig. 1b-d)**. We thus speculate that enough Fe(II) can be coordinated in the catalytic center of PCOs even at very low intracellular Fe concentrations, both in preformed enzymes, which are not likely to lose their cofactor, and in newly synthesized ones.

Among the numerous Cys2 proteins encoded by the Arabidopsis proteome, four group Ib bHLH factors can be found, namely bHLH038, 039, 100 and 101. Recently, (Kozlic et al., 2022) detected the marginal stabilization of an (Ala2)bHLH038-GFP protein expressed in yeast. Here, using a heterologous yeast strategy (Puerta et al., 2019) and transient protoplast transformation, we found that bHLH039 is not a susceptible substrate of the Cys N-degron pathway **(Fig. 2)**. Considering our conclusions on PCO prioritization in terms of Fe usage, we speculate that subtracting the Ib bHLHs from the Cys-NDP is functional to ensure their accumulation under mild Fe-deficient conditions. Our bHLH039-Fluc constructs displayed very low stability, according to the luminescent output, although perfectly detectable bHLH039-mCherry levels have been reported by (Trofimov et al., 2019)in Arabidopsis leaves. The data hint at the presence of some uncharacterized degradation signal in the N-terminal part of the protein, associated with bHLH039 turnover both in yeast and plant leaf protoplasts, in Fe-replete conditions.

We investigated the implications of PCO inhibition under severe Fe-deficiency in light of ERFVII regulation by co-occurring environmental factors, in addition to oxygen availability. We recorded the transcriptional signature of hypoxic responses in wild type Arabidopsis plants growing in chronic Fe deficiency conditions, indicating that Fe starvation can affect hypoxia signaling depending on its severity. Moderate Fe stress did not invoke hypoxic signaling **(Fig. 3**, **Fig. 5, Supplemental Fig. 3)**, while severe chronic Fe deficiency did **(Fig. 4)**. In axenic seedlings, stabilization of ERFVII factors and hypoxic gene induction in chronic Fe deficiency conditions has been reported in previous studies (Carbonare et al., 2019; Zubrycka et al., 2023). Here, notably, we observed the same response both in seedlings cultivated on synthetic agar media and in adult plants growing on soil, suggesting that the stimulation of hypoxic signaling is part of a general strategy under Fe insufficiency. We cannot rule out that altered oxygen consumption might take part to this phenomenon. Mitochondrial electron transport chain (mETC) complexes are highly dependent on Fe availability and therefore it has been demonstrated that severe Fe deficiencies strongly affect mitochondrial functionality both at root and leaf level. Despite the impairment of respiration, increased oxygen consumption has been indeed reported in cucumber roots subjected to severe Fe-deficiency (Vigani et al., 2009). This effect has been attributed to the increase of other O_2_-consuming processes, such as Fe^3+^-chelate reductase and some ROS detoxification activities.

The comparison of wild type and *erfVII* pentuple mutant plants (Abbas et al., 2015) shed light on the contribution of ERFVII stabilization to Fe deficiency responses. The down-regulation of some of the Fe-starvation markers observed in mutant seedlings **(Fig. 4c)** and rosette tissues **(Fig. 3a)**, particularly *bHLH038* or *039*, suggests that the ERFVII may contribute to the acclimation to chronic Fe deprivation by acting as positive regulators of a subset of starvation-responsive genes. The *erfVII* mutation did not affect growth on Fe- substrates **(Fig. 3b and 4a-b)**, but it led to some alterations in micronutrient content at the seedling stage **(Fig. 4d)**. Moreover, combination of submergence with alkaline soil highlighted a positive role for the ERFVII factors in nutrient mobilization to leaves, particularly micronutrients **(Fig. 6)**. Our experiments expand the role of ERFVIIs to nutrient management under submergence and dampening of starvation responses in the leaves of plants experiencing moderate chronic iron deficiency **(Fig. 7)**.

A well-known player in submergence responses is the phytohormone ethylene, which in submerged tissues accumulate rapidly due to a combination of enhanced biosynthesis and restrained outward diffusion (Sasidharan et al., 2018; Sasidharan & Voesenek, 2015; Voesenek et al., 1993). On the other hand, transcriptome analyses and genetic evidence indicate that ethylene is also involved in the promotion of Fe uptake in plants through the regulation of FIT (García et al., 2014;Lingam et al., 2011). Ethylene is thus considered to be a positive regulator of Fe deficiency responses, although full understanding of the underlying mechanisms is yet to be gained (Li & Lan, 2017; Lucena et al., 2015). Submergence induced ethylene build-up has been shown to contribute to the ERFVII-mediated induction of anaerobic gene expression indirectly, through phytoglobin-mediated NO scavenging and consequent ERFVII stabilization (Hartman et al., 2019). This mechanism explains the observation that ethylene pretreatment can prime ERFVII-mediated responses to subsequent hypoxia, in which ethylene entrapment is not expected to take place. We observed a down-regulation of the transcriptional Fe- markers by prolonged hypoxia **(Fig. 5)**, which was also present in submerged leaves of Fe-starved plants **(Fig. 7)**. The effect might be due to a global strategy of selective transcriptional prioritization under hypoxia. However, this finding is compatible with previous reports of a specific function of the ERFVII family factors RAP2.3 and RAP2.12 as negative regulators of genes involved in Fe uptake under Fe deficiency conditions (Liu et al., 2017). Future studies will help disentangle the contribution of ethylene- and ERFVII-mediated signalling to nutrient acquisition under combined nutritional and submergence stresses.

It is accepted that flooding tolerance should include traits associated with the alleviation of nutrient deficiencies and phytotoxicity issues that can both arise after prolonged submergence or soil waterlogging (Yuan et al., 2023). Waterlogged soils undergo complex modifications of pH and microbial activities, as a consequence of redox changes, that, depending on the specific edaphic properties, can often lead to the increase of nutrient availability above the toxicity threshold (typically, Zn(II), Mn, Fe(II), or S) (Shabala, 2011; Tamang & Fukao, 2015). Fe toxicity, for instance, is recognized as a relevant problem in several rice cropping areas of the world (Mahender et al., 2019; Shabala, 2011). On the other hand, submergence is expected to impair nutrient acquisition as a consequence of limited transpiration and inhibition of active membrane transport. Reduced mobilization of nutrients to the shoots has been indeed frequently noticed in waterlogged cereals (Colmer & Greenway, 2011; Singh & Setter, 2017; Steffens et al., 2005). Our finding that plant ERFVII factors participate to nutrient management in plants growing on alkaline substrate might be of potential interest for the improvement of crop responses to waterlogging/submergence events in calcareous and Fe-deficient soils, which are believed to account to 30% of soils globally (Mahender et al., 2019).

In conclusion, this study provided novel insights into the interactions between low Fe- and low O_2_-response pathways in plants and their implications for plant stress responses. Understanding the intricate regulatory networks that govern Fe homeostasis and O_2_ sensing is crucial for enhancing plant growth, improving stress tolerance, and developing sustainable agricultural practices to withstand the current environmental challenges. Further investigations will be needed to gain a more detailed view of ERFVII role in fine-tuning gene expression during chronic Fe deficiencies and their interplay with ethylene, during combined Fe and submergence stresses.

## Author Contributions

Y.T., Conceptualization, Methodology, Writing-Original Draft, Investigation, Visualization. M.M., Methodology, Investigation, Visualization. M.L. and N.L.M., Methodology, Investigation. G.A., A.S. and S.D., Investigation. P.P., Conceptualization, Supervision, Writing - Review & Editing, Funding acquisition. B.G. and G.V., Conceptualization, Methodology, Writing-Original Draft, Visualization, Supervision, Funding acquisition, Project administration.

## Acknowledgements

We thank Dr Monirul Islam for technical assistance in the pot experiments.

