## Supplemental Figures and Tables for "Severe Fe deficiency promotes hypoxia inducible responses in *Arabidopsis thaliana*"

### Supplemental Information

SI Figures 1-3

SI Tables 1-3

### SI Figures

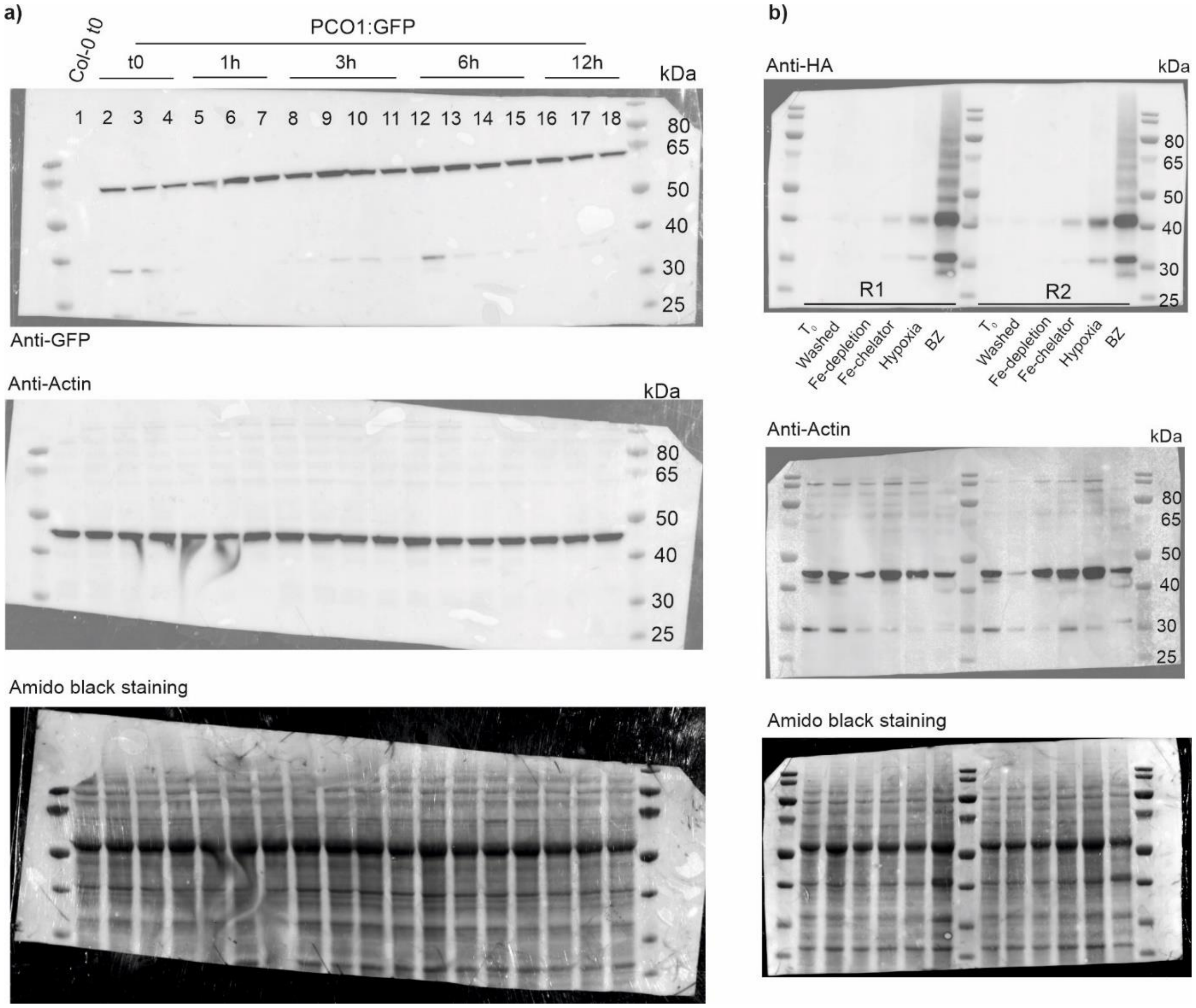


**Supplemental Figure 1. Supporting blot images. (a)** Full blots from Figure 1d. Lanes 1-2, 7-8 and 15-16 are displayed in the main text. **(b)** Full blots from Figure 1f. Two biological replicates were performed.

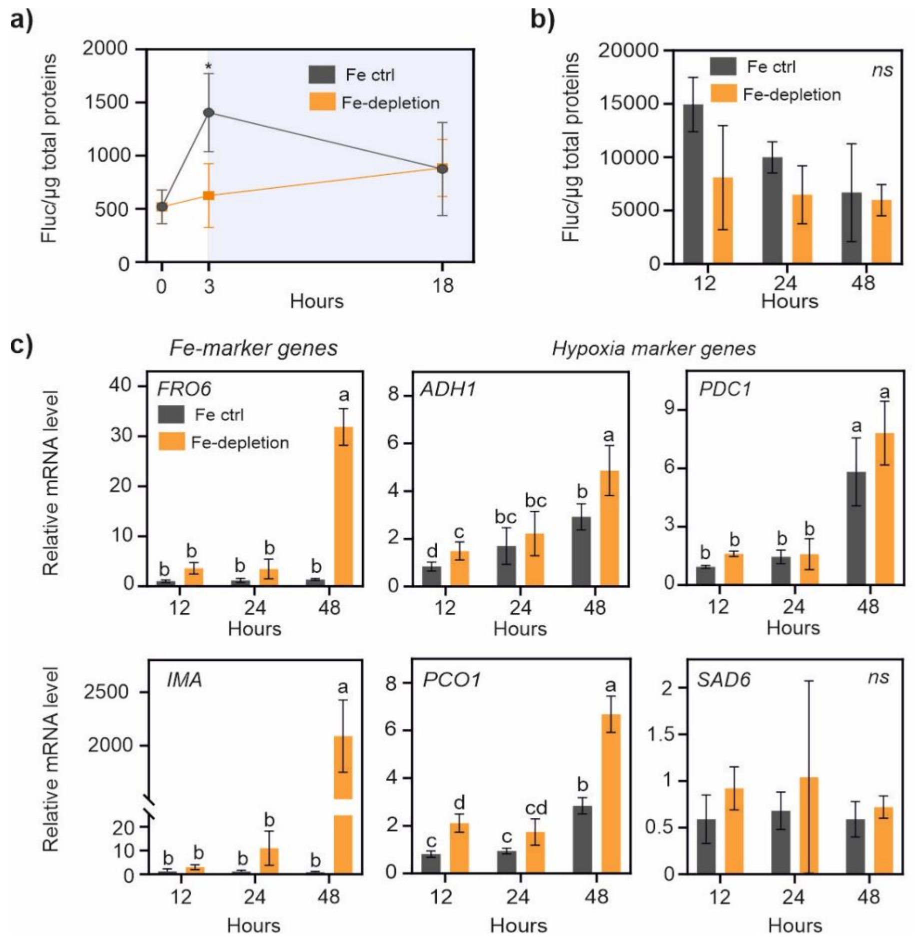


**Supplemental Figure 2. Effects of prolonged Fe-depletion on RAP2.12. (a)** RAP2.12 stability in 7 day-old *28RAP2.12Fluc* seedlings after apoplast washing, followed by shifting to fresh Fe-deficiency medium and maintenance under continuous darkness for 3 or 18 h. Data are mean ± SD (n=5). **(b)** 28RAPFluc stability after shifting of seedling to fresh iron-deficiency medium under neutral photoperiod. $t_0$ corresponded to the end of day (8 PM). All data are mean ± SD (n=5). **(c)** Expression (mean ± SD, n=5) of marker genes in the Col-0 ecotype, treated as indicated in (b). Asterisks mark statistically significant differences between control and treated samples after Student's t-test at each time point (P<0.05).

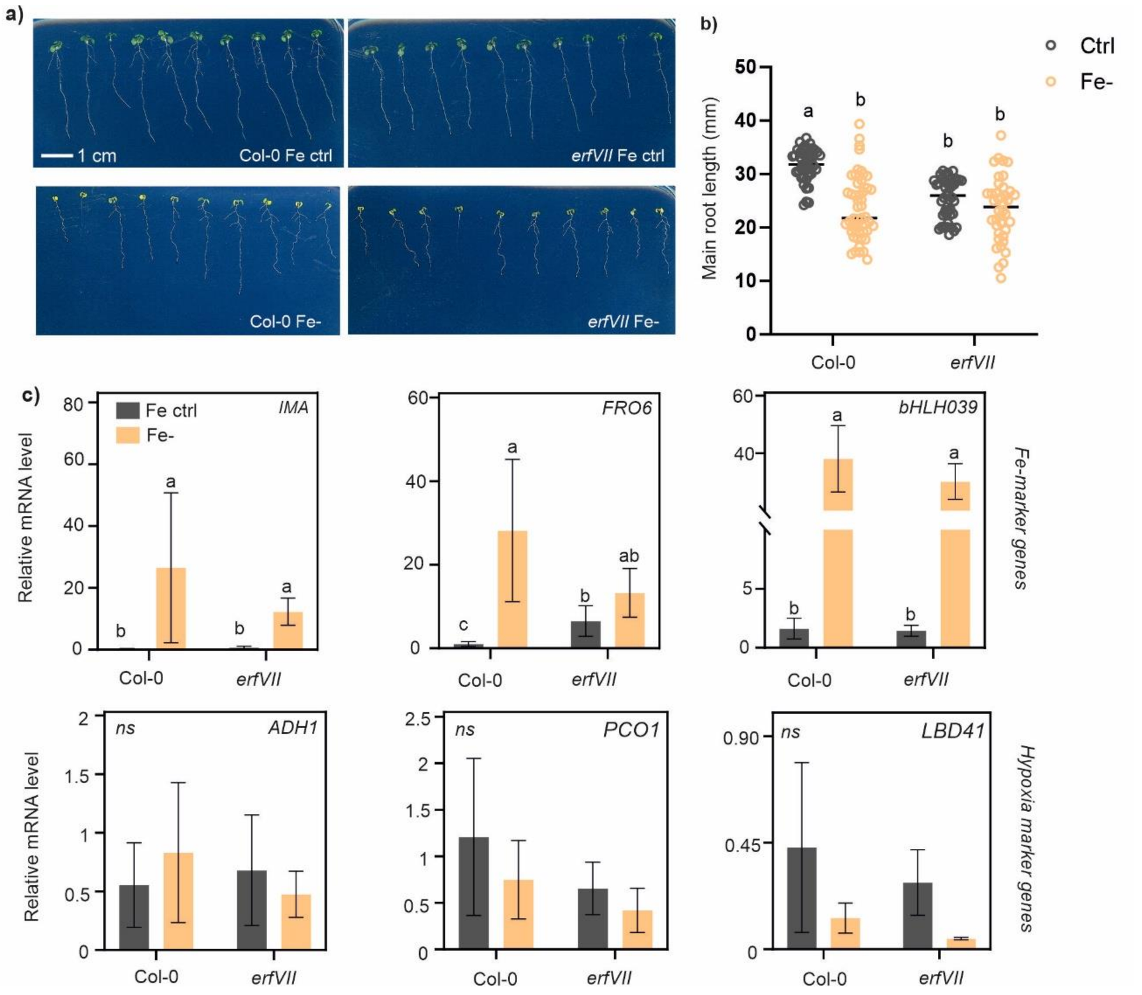


**Supplemental Figure 3. Phenotypic and molecular responses in *erfVII* mutants under moderate chronic iron-deficiency. (a)** Representative pictures of wild-type and *erfVII* seedlings grown for 10 days on control or Fe- plates, in presence of 0.9% agar and 1% sucrose (w/v). Five replicate plates were observed for each experimental thesis. **(b)** Quantification of primary root length in the plantlets depicted in (a). Data are mean ± SD (n=57). **(c)** Expression of iron starvation and anaerobic markers in the same experiment. Expression values (mean ± SD, n=5) are presented as normalized to a wild-type control sample. Distinct letters indicate statistically significant differences after two-way ANOVA and Tukey-Kramer post-hoc test ($P < 0.05$).

### SI Tables

**Supplemental Table 1. List of qPCR primers used in this study.**

| Gene name | AGI code | Forward primer sequence | Reverse primer sequence |
|---|---|---|---|
| *BHLH038* | *AT3G56970* | TCAACGGTTTCTGCCACTAG | ACATCCACAAGAACAAACCCA |
| *BHLH039* | *AT3G56980* | TGTTTCTGTTTCGTCGGAGG | TAATTTTCCTGCGACGGTCA |
| *FRO6* | *AT1G01580* | GCTTCCGCCGATTTCTTAAGGC | AACGGAGTTATCCCGCTTCCTC |
| *IMA* | *AT1G47400* | GGCCATCAAGAGATTTGACCATGC | TGCCACTCGAGAACTATCTACCAC |
| *ADH1* | *AT1G77120* | TATTCGATGCAAAGCTGCTGTG | CGAACTTCGTGTTTCTGCGGT |
| *HB1* | *AT2G16060* | TTTGAGGTGGCCAAGTATGCA | TGATCATAAGCCTGACCCCAA |
| *HRA1* | *AT3G10040* | ACAACCACCGCAACAGAATCC | TCTCCGCAATTCTCGCCAT |
| *LBD41* | *AT3G02550* | TGAAGCGCAAGCTAACGCA | ATCCCAGGACGAAGGTGATTG |
| *PCO1* | *AT5G15120* | ATTGGGTGGTTGATGCTCCAATG | ATGCATGTTCCCGCCATCTTC |
| *SAD6* | *AT1G43800* | TTGGCAACCCGCTTCTTTCTTACC | TTTCCCTCAGCTCACGAACCTG |
| *UBQ10* | *AT4G05320* | GGCCTTGTATAATCCCTGATGAATAAG | AAAGAGATAACAGGAACGGAAACATAGT |

**Supplemental Table 2. Ion quantification in seedlings by ICP-MS.** Absolute concentration of mineral ions (µg $g^{-1}$ dry weight) from n=3 replicates. Raw data supporting Figure 4 in the main text.

| | **Col-0 Fe ctrl** | **Col-0 Fe-** | ***erfvii* Fe ctrl** | ***erfvii* Fe-** |
|---|---|---|---|---|
| Na | 3571.45±1746.57 | 3440.16±1103.19 | 3341.4±2156.2 | 6822.91±1656.57 |
| Mg | 1834.33±280.24 | 2270.85±361.91 | 2291.29±217.01 | 3096.55±149.61 |
| K | 44497.57±7208.97 | 55482.42±11273.34 | 47032.85±3622.96 | 57659.21±5181.58 |
| Ca | 4731.82±833.24 | 6068.61±852.65 | 5791.48±541.23 | 8274.12±970.8 |
| Mn | 133.51±21.06 | 368.48±18.1 | 137.27±11.16 | 532.52±66.38 |
| Fe | 229.46±59.31 | 94.89±14.1 | 194.59±38.4 | 164.4±8.83 |
| Co | 1.32±0.29 | 23.58±5.04 | 1.44±0.36 | 23.07±5.67 |
| Cu | 13.63±0.56 | 12±5.32 | 5.61±1.58 | 17.1±2.2 |
| Zn | 198.61±39.79 | 1000.21±38.46 | 154.87±38.7 | 992.63±145.02 |
| Mo | 41.84±1.23 | 44.46±3.75 | 11.19±2.32 | 132.34±16.69 |

**Supplemental Table 3. Ion quantification in rosette leaves by ICP-MS.** Absolute concentration of mineral ions (µg $g^{-1}$ dry weight) from n=3 replicates. Raw data supporting Figure 6 in the main text. K in submergence control samples was present above the detection limit of the instrument and could not be determined.

| | | **Col-0** | | |
|---|---|---|---|---|
| **Ion** | **Normoxia ctrl** | **Normoxia CaO** | **Submergence ctrl** | **Submergence CaO** |
| Na | 1631.17 ± 126.76 | 2716.72 ± 91.85 | 1562.13 ± 403.16 | 3546.48 ± 686.34 |
| Mg | 11182.46 ± 1653.93 | 7066.47 ± 147.86 | 8683.28 ± 1343.22 | 8721.52 ± 1718.04 |
| K | 47933.4 ± 7319.26 | 39483.85 ± 4026.18 | over | 44879.21 ± 9150.63 |
| Ca | 44684.27 ± 7394.68 | 43688.52 ± 1911.98 | 32861.91 ± 4844.67 | 54890.25 ± 10168.33 |
| Mn | 96.85 ± 68.51 | 140.26 ± 15.56 | 44.1 ± 6.68 | 266.08 ± 137.24 |
| Fe | 205.44 ± 12.44 | 196.98 ± 5.29 | 194.99 ± 23.7 | 300.02 ± 53.68 |
| Cu | 9.9 ± 1.01 | 12.01 ± 3.67 | 8.69 ± 0.91 | 14.58 ± 2.57 |
| Zn | 141.12 ± 40.68 | 118.92 ± 8.12 | 290.69 ± 214.11 | 437.45 ± 459.88 |
| Mo | 18.33 ± 4.14 | 24 ± 5.99 | 14.53 ± 2.92 | 33.9 ± 12.55 |
| P | 11.63 ± 1.59 | 11.24 ± 0.62 | 10.54 ± 1.34 | 14.54 ± 3.16 |
| | | ***erfvii*** | | |
| **Ion** | **Normoxia ctrl** | **Normoxia CaO** | **Submergence ctrl** | **Submergence CaO** |
| Na | 1594.22 ± 373.6 | 2667.76 ± 549.28 | 1825.78 ± 415.66 | 2152.56 ± 82.14 |
| Mg | 11355.15 ± 1435.02 | 7709.15 ± 1233.38 | 8721.03 ± 1519.49 | 6416.53 ± 1356.52 |
| K | 41930.21 ± 3325.43 | 34210.44 ± 7012.78 | 38960.89 ± 2412.41 | 30025.99 ± 1361.63 |
| Ca | 43292.64 ± 4745.24 | 48787.99 ± 8624.49 | 37522.87 ± 1434.49 | 39229.35 ± 9194.66 |
| Mn | 59.45 ± 1.59 | 212.99 ± 112.04 | 68.42 ± 24.91 | 105.82 ± 36.66 |
| Fe | 224.29 ± 35.31 | 234.44 ± 3.24 | 233.99 ± 22.19 | 208.82 ± 18.27 |
| Cu | 9.62 ± 0.52 | 10.46 ± 2.19 | 11.27 ± 0.92 | 8.37 ± 1.68 |
| Zn | 87.8 ± 4.94 | 87.07 ± 5.28 | 153.04 ± 39.79 | 106.42 ± 17.24 |
| Mo | 15.47 ± 1.71 | 25.13 ± 11.22 | 21.01 ± 8.78 | 20.98 ± 2.39 |
| P | 11.99 ± 0.62 | 10.43 ± 2.53 | 11.8 ± 0.53 | 10.04 ± 1.68 |
